# MpNPR modulates lineage-specific oil body development and defence against gastropod herbivory in *Marchantia polymorpha*

**DOI:** 10.1101/2025.11.17.688000

**Authors:** Loreto Espinosa-Cores, Santiago Michavila, Marina Gonzalez-Zuloaga, Roberto Solano, Selena Gimenez-Ibanez

## Abstract

The transition to land exposed early plants to novel biotic constraints, creating strong selection pressures that drove the evolution of the plant immune system. Extant land plants comprise two major groups, tracheophytes and bryophytes, which include liverworts, hornworts and mosses. These two groups diverged early after land colonization, and thus, immune mechanisms have also been subjected to independent evolution. Here, we investigated the function of the unique NPR in the liverwort *Marchantia polymorpha* and report that it exhibits a specialised immune function relative to its orthologs in tracheophytes. We show that MpNPR modulates the formation of oil bodies, a synapomorphy of liverworts, which function as storage organelles for secondary metabolites and defence against arthropods. We found that MpNPR interacts with MpERF13, a transcription factor controlling oil body differentiation, and that plants with altered levels of MpNPR are misregulated in the expression of MpERF13-dependent genes. Furthermore, we uncover that MpNPR and MpERF13 are required for the defence against snail herbivory. Finally, we show that the ability of MpNPR to induce oil body formation and resistance to gastropods is fully dependent on MpERF13. Taken together, our results demonstrate that NPR mediates a novel regulatory pathway in lineage-specific oil body formation and immunity against gastropod herbivory through the MpERF13 TF in *Marchantia,* while also pinpoint a novel specialised evolutionary trajectory of NPR in liverworts.

## INTRODUCTION

The colonisation of the terrestrial environment by plants was a key step in evolution that paved the way for all terrestrial life. On land, early plants were rapidly exposed to a multitude of novel biotic stresses, constituting strong selection pressures that acted as a driving force for the evolution of new sophisticated defence mechanisms to quickly adapt or perish ^1,2^. This included the emergence of effective immune responses against a large diversity of microbes with contrasting infective strategies and lifestyles such as bacteria, fungi, oomycetes and herbivores, among many others. Extant land plants comprise two major groups, tracheophytes (vascular plants) and bryophytes (non-vascular plants). Tracheophytes contain four plant lineages (lycophytes, ferns, gymnosperms and angiosperms), while bryophytes include the three lineages liverworts, hornworts and mosses ^3–5^. These two groups diverged from each other early after land colonization, and thus, immune mechanisms against invaders have also been subjected to independent evolution over the past 400 million years ^5^. This might have led to a large diversity of immune mechanisms, which encompasses well-conserved across land plants’ defence systems, such as cell surface immune receptors and metabolic routes, and also, lineage-specific innovations for immunity ^2^. Lineage-specific plant immunity might be the result of singular co-evolution with pathogen pressures, and represent diversified reservoirs of immune mechanisms that are exclusive to certain plants.

Most liverworts contain oil bodies within their cells, which represents a typical synapomorphy of this lineage, and a likely liverwort-specific innovation for immunity against herbivores ^6–9^. Oil bodies are intracellular specialized organelles that synthesize and store a large diversity of bioactive cytotoxic metabolites, such as terpenoids or bisbibenzyls, as major chemical constituents ^6,8,10–13^. Their specific characteristics, such as size, shape, and colour, vary between plant species ^8^. Some liverworts, like in the *Radula* genus contain oil bodies in almost every cell, while in others, such as *Marchantia polymorpha* (*Marchantia*), oil bodies are large and differentiate only in specialized cells referred to as idioblasts in a relative constant proportion ^14^. Recent studies in *Marchantia* have provided significant insights into the mechanisms governing oil body cell differentiation and formation, highlighting key transcription factors (TFs), such as MpERF13, MpC1HDZ, MpTGA, and MpMYB02 in this process ^6,7,15,16^. Among all, MpERF13 and MpC1HDZ are master TFs required for oil body cell differentiation and formation that act likely by regulating the expression of MpSYB12B, an SNARE protein that coordinates the redirection of the secretory pathway towards the formation of the oil body ^6,7^. In contrast, MpMYB02 likely acts later in the development of the oil body cell and is also involved in the biosynthesis of secondary metabolites such as marchantins that are stored at the organelle ^10,15,17^. Moreover, retention of additionally synthetised sesquiterpenes within the oil body further require the Mp*ABCG1* transporter ^12^. Notably, recent targeted functional analyses in *Marchantia* have shown that mutants for the Mp*ERF13* and Mp*C1HDZ* TFs, which are depleted of oil body cells, are also more susceptible to the attack of the terrestrial isopod *Armadillidium vulgare* ^6,7^. Conversely, gain-of-function mutants of Mp*ERF13* or genome-edited mutant lines of MpTGA, which show a significant increase in oil body numbers, are more resistant to feeding by this arthropod ^7,16^. These results have stablished the first evidences for the direct role of oil bodies in controlling herbivory by arthropods in *Marchantia*. However, oil bodies are likely to be not only effective in protecting against the attack of arthropods as early experiments performed more than a century ago already associated the presence of these organelles to the feeding behaviour of land snails ^18^. While all this recent work has paved the way for starting to understand the regulatory circuits controlling oil body cell differentiation and function, which host regulators control TF-driven oil body formation and liverwort-specific immunity in *Marchantia* remains enigmatic.

In angiosperms, *NPR* (*non-expressors of PR*) genes such as *NPR1, NPR3*, and *NPR4* have been extensively studied for their essential roles as hormonal salicylic acid (SA) sensors and executors of SA-dependent plant defences against biotrophic and hemibiotrophic pathogens, although with different binding affinities for the hormone and immune outcomes ^19–23^. While NPR1 is a positive regulator of SA signalling and related defence responses, NPR3/NPR4 act as repressors ^21^. These NPRs features represent a commonality for multiple plants from angiosperms ^24^. *Marchantia* has a unique MpNPR ortholog, which is a substantial difference compared to the *Arabidopsis* genome ^25^. Its recent characterization has revealed that Mp*NPR* can complement the At*npr1* mutant, but not At*npr3npr4,* when ectopically expressed in *Arabidopsis* ^25,26^. This goes in line with that previously observed for a PpNPR from the moss *Physcomitrium patens* ^27^. Paradoxically, mutations in Mp*NPR* gene do not result in At*npr1*-like phenotypes in *Marchantia*. In fact, Mp*npr* mutants show enhanced resistance to the hemibiotrophic bacterium *Pseudomonas syringae*, hypersensitivity to SA exogenous treatments and notably, unaltered SA-induced transcriptional reprograming compared to wild-type *Marchantia* plants ^25^. These unexpected results indicate that Mp*NPR* may not be the master regulator of SA-induced responses in this liverwort. In support of this, hornworts lack NPR genes but still produce SA for still unknown purposes ^28–30^. Additionally, recent analysis in Mp*npr* mutants have identified roles for MpNPR in the regulation of abiotic stresses, including thermomorphogenesis and far-red light responses, and also developmental processes such as sexual reproduction ^16,25^. This has led to the hypothesis that, despite some NPR biochemical properties might be partially conserved between bryophytes and tracheophytes, MpNPR might have been initially acquired by liverworts to different purposes than vascular plants ^2,25^. Taken together, the recent data suggests that NPR-associated pathways might have evolved distinctly in divergent land plant lineages, while also opens the intriguing question of whether NPR, which is considered a master regulator on plant defences in most plants, also contribute to immunity in bryophytes to any extent.

Here, we investigated the role of MpNPR and report that it exhibits a specialised immune function unique to liverworts. Our results show that MpNPR controls lineage-specific oil body formation and defence against snail herbivory through the master MpERF13 TF in *Marchantia,* further indicanting that the defensive role of oil bodies is not restricted to arthropods and that might be general against herbivore attack. Globally, our suggests that NPRs from liverworts were co-opted towards the control of the oil body to fend off against the selective pressures that likely early herbivores impose on ancestral liverworts.

## RESULTS

### MpNPR is a positive regulator of oil body formation

To gain insight into the NPR function in *Marchantia*, we generated Mp*npr* mutants by conventional CRISPR/Cas9 technology in the WT Tak-1 background ^31^. We obtained two different alleles, Mp*npr-3^ge^* and Mp*npr-4^ge^*, which contained a 22 bp deletion and a 37 bp insertion in the first exon of Mp*NPR*, respectively (Figure S1A). Both alleles resulted in a premature stop codon and, therefore, non-functional MpNPR peptides lacking all fundamental domains for an NPR protein (Figure S1A). Our Mp*npr-3^ge^* and Mp*npr-4^ge^*alleles did not show obvious developmental defects (Figure S1B), as alleles described previously ^16,25^. To understand the biological role of MpNPR, we performed a detailed phenotypic characterization of the mutant alleles. It has been previously described that the thallus of *M. polymorpha* grown on axenic conditions contain a relative low number of oil bodies, while specific environmental conditions such as starvation and non-axenic cultivation increase the accumulation of these organelles ^6,10^. Thus, we analysed the formation of oil bodies in gemmae from Tak-1 and Mp*npr* mutants grown on axenic and non-axenic conditions using vermiculite as substrate. As observed before for oil body cells in the thallus of *M. polymorpha* ^6,10^, gemmae from non-axenically grown Tak-1 plants showed significantly higher amounts of oil bodies, and specially, decreased dispersion in the number of these organelles among gemmae, compared to plants grown on axenic conditions (Figure S2). Under non-axenic conditions, gemmae from both Mp*npr* mutant alleles showed a significant reduction in the number of oil bodies compared to Tak-1 plants (Figure 1A and 1B). In contrast, under axenic conditions, gemmae from Mp*npr* mutants grown *in vitro* did not display significant differences compared to Tak-1 (Figure S3), as it has been previously described ^16^. This indicates that MpNPR is not essential for basal levels of oil bodies, but contributes to the formation of these organelles during environmental-induced modulation of oil body development.

**Figure 1.**
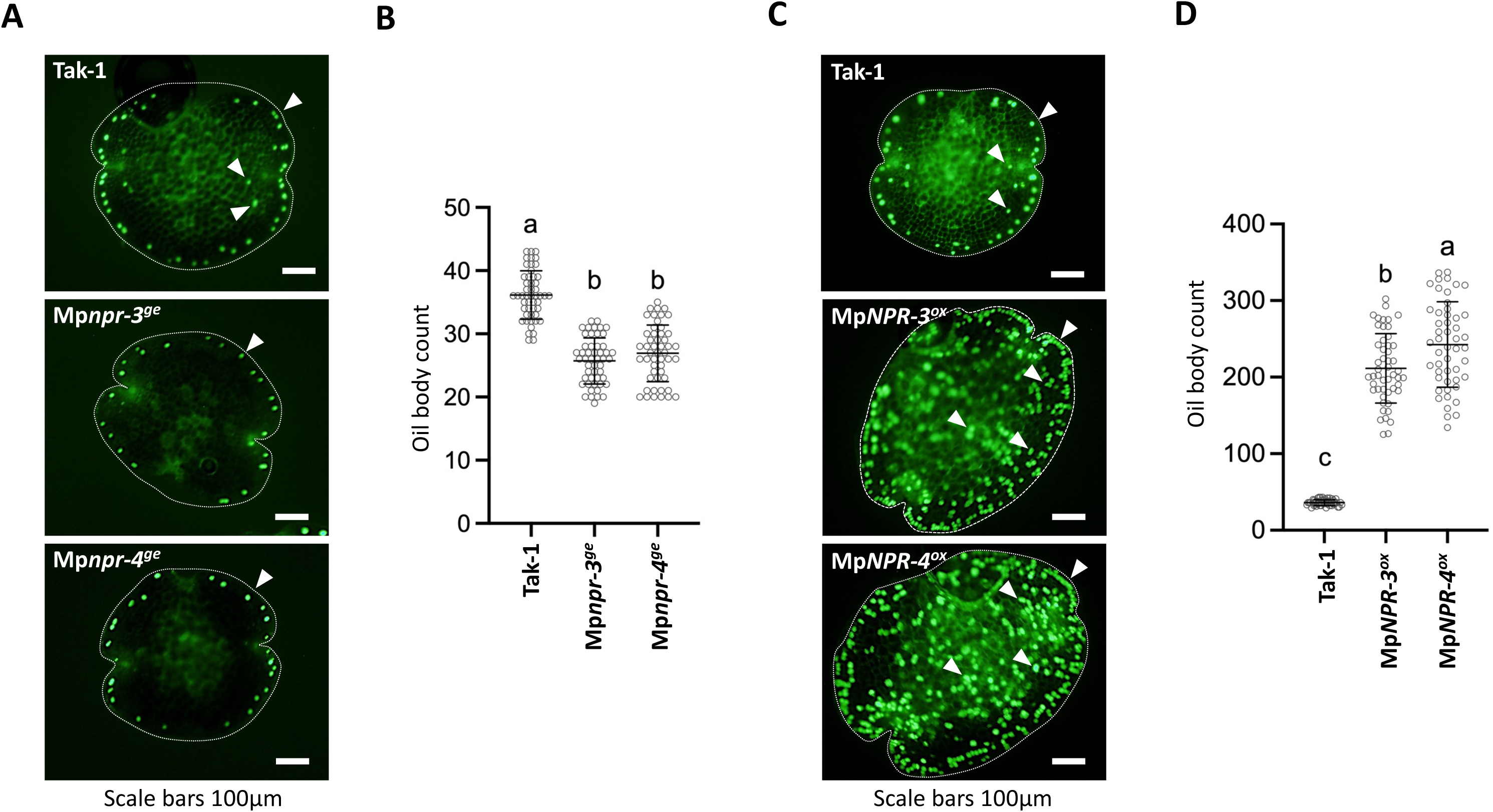
MpNPR is a positive regulator of oil body formation. **(A)** Phenotype of BODIPY-stained 0-d-old gemmae from WT Tak-1 and Mp*npr* mutant plants under fluorescence microscopy. Oil bodies are indicated by arrows. **(B)** Number of oil bodies visualized by BODIPY-staining in 0-d-old gemmae of WT Tak-1 and Mp*npr* mutant plants. Bars indicate means ± SD (n = 50). Letters indicate statistically significant groups (one-way ANOVA, Tukey’s HSD, *p* < 0.01). **(C)** Phenotype of BODIPY-stained 0-d-old gemmae from WT Tak-1 and Mp*NPR* overexpressing plants under fluorescence microscopy. Oil bodies are indicated by arrows. **(D)** Number of oil bodies visualized by BODIPY-staining in 0-d-old gemmae of WT Tak-1 and Mp*NPR* overexpressing plants. Bars indicate means ± SD (n = 50). Letters indicate statistically significant groups (one-way ANOVA, Tukey’s HSD, *p* < 0.01).

To further study the function of MpNPR during the development of oil bodies, we next evaluated the impact of increasing endogenous MpNPR levels in *Marchantia.* To do this, we generated transgenic *Marchantia* plants overexpressing Mp*NPR-citrine* under the control of the Mp*EF1⍺* promoter in the Tak-1 background. Western blotting assays confirmed protein accumulation of MpNPR-citrine (Figure S4A). Compared to Tak-1, ectopic Mp*NPR* expression reduced thallus growth while maintaining the typical dichotomous branching from apical meristems of this liverwort (Figure S4B). However, the most outstanding phenotype of *Marchantia* plants overexpressing Mp*NPR* (Mp*NPR-3^ox^* and Mp*NPR-4^ox^*) was a striking overaccumulation in the number of oil bodies in gemmae (Figure 1C and 1D). Whereas WT Tak-1 gemmae formed around 40 oil bodies predominantly accumulating near the gemmae margin, gemmae overexpressing Mp*NPR* contained an average above 200 oil body cells, distributed throughout all the gemmae tissue (Figure 1C and 1D). These data indicate that Mp*NPR* is a positive regulator of oil body formation in *Marchantia*.

### MpNPR interacts with MpERF13 transcription factor

Recent studies have provided insights into the molecular mechanisms controlling oil body formation in *Marchantia*, uncovering key TFs involved in oil body cell differentiation and maturation ^8^. These include TFs from several families, such as MpERF13 ^7^, MpC1HDZ ^6^ and MpMYB02 ^15^, as positive regulators of oil body cell differentiation and formation. Conversely, the MpTGA was identified as a negative regulator for the formation of these structures ^16^. Among all, the MpERF13 TF from the AP2-ERF family called our attention as a recently described gain-of-function allele of Mp*ERF13* (Mp*erf13^GOF^*) showed a significant increase in oil body numbers reminiscent to what we observed for our overexpressing Mp*NPR* gemmae ^7^. Thus, we directly compared oil body numbers in gemmae from Mp*erf13^GOF^*, and also an already characterized genome-edited mutant line of Mp*ERF13* (Mp*erf13-1^ge^*), with mutant and overexpressing lines of Mp*NPR* (Mp*npr-4^ge^* and Mp*NPR-4^ox^*) ^7^. As previously described ^7^, Mp*erf13-1^ge^* gemmae were completely depleted of oil bodies while Mp*erf13^GOF^*gemmae significantly over accumulated these organelles along all the gemma compared to Tak-1 (Figure 2A). Mp*npr* gemmae showed significantly lower amounts of oil bodies compared to Tak-1, but far from Mp*erf13-1^ge^* (Figure 2A and 2B). In contrast, gemmae overexpressing Mp*NPR* showed increased oil body numbers and a distribution pattern for these specialized structures rather similar to that observed for Mp*erf13^GOF^* (Figure 2A and 2C). While these results indicate that MpNPR is not the unique protein modulating oil body formation in *Marchantia*, the similarities between Mp*NPR^ox^* and Mp*erf13^GOF^* raise the possibility that MpNPR could be connected to the MpERF13 TF to modulate oil body cell differentiation in *Marchantia*.

**Figure 2.**
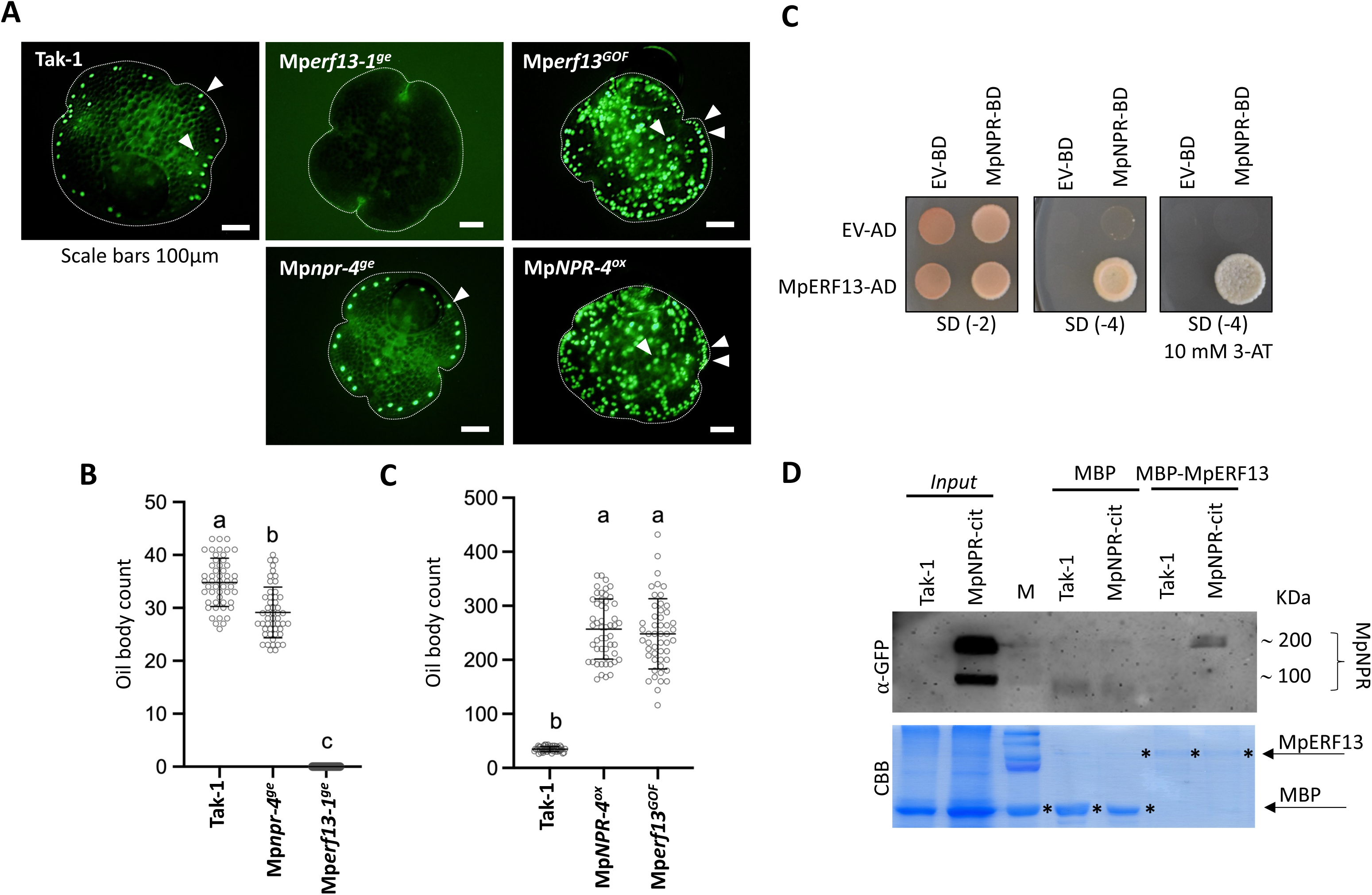
MpNPR interacts with MpERF13 transcription factor. **(A)** Phenotype of BODIPY-stained 0-d-old gemmae from WT Tak-1, Mp*erf13-1^ge^,* Mp*erf13^GOF^,* Mp*npr-4^ge^* and Mp*NPR-4^ox^* plants under fluorescence microscopy. Oil bodies are indicated by arrows. **(B)** Number of oil bodies visualized by BODIPY-staining in 0-d-old gemmae of WT Tak-1, Mp*erf13-1^ge^* and Mp*npr-4^ge^* mutant plants. Bars indicate means ± SD (n = 50). Letters indicate statistically significant groups (one-way ANOVA, Tukey’s HSD, *p* < 0.01). **(C)** Number of oil bodies visualized by BODIPY-staining in 0-d-old gemmae of WT Tak-1, Mp*erf13^GOF^* and Mp*NPR-4^ox^* plants. Bars indicate means ± SD (n = 50). Letters indicate statistically significant groups (one-way ANOVA, Tukey’s HSD, *p* < 0.01). **(D)** Yeast two-hybrid (Y2H) interaction assays between MpNPR and MpERF13. BD, binding domain (pGBKT7 vector); AD, activation domain (pGADT7 vector). Growth on plasmid-selective media SD (-2) (minimal media lacking leucine (L) and tryptophan (T)), and interaction-selective media SD (-4) (lacking leucine (L), tryptophan (T), adenine (A) and histidine (H), in the presence or absence of 10 mM of 3-AT) are shown. **(D)** MpNPR interacts with MpERF13 TF in pull-down (PD) assays. Immunoblots with anti-GFP antibody of MpNPR-citrine recovered after PD experiments using crude protein extracts from Mp*NPR-4^ox^* (*pro*Mp*EF1α:*Mp*NPR-citrine*) or Tak-1 plants, and resin-bound recombinant MBP or MBP-fused MpERF13 protein. Input lines show the level of expression of MpNPR protein in transgenic and control plants. CBB staining shows the amount of recombinant MpERF13-MBP or MBP proteins used in the resin (bottom).

In angiosperms, NPR proteins directly interact with several different TFs and TF families to regulate their responses ^32–37^. In addition, further confocal microscopy on our overexpressing MpNPR-citrine lines showed that this protein is nuclear localized in stable *Marchantia* transgenic plants (Figure S4C), supporting the previous observed nuclear localization of MpNPR when transiently expressed in heterologous systems ^16,26^. We reasoned that similarities in the content of oil bodies between Mp*NPR^ox^* and Mp*erf13^GOF^*plants imply that MpNPR might target MpERF13 directly. To test this, we examined the ability of MpNPR to interact with the MpERF13 in a yeast two-hybrid assay (Y2H). MpNPR interacted with MpERF13 in these experiments (Figure 2C). To further confirm the MpNPR-MpERF13 association, we expressed the MpERF13 fused to MBP in *E. coli* and purified the protein for pull-down (PD) analysis with cell extracts of transgenic plants overexpressing MpNPR-citrine. The MpNPR–MpERF13 interaction was also confirmed by PD assays (Figure 2D). Noteworthy, MpNPR was detected as a double band in these PD experiments, which might correspond to monomeric and homodimer forms of MpNPR-citrine (∼ 95 kDa), as described for the *Arabidopsis* NPR1 ortholog ^20^ (Figure 2D). Thus, our results indicate that MpERF13 TF is a likely target of MpNPR in *Marchantia*.

### MpNPR regulates the expression of MpERF13-dependent genes

Combined RNAseq-based transcriptomic profiling of Mp*erf13-1^ge^* and Mp*erf13^GOF^* mutant plants have uncovered core genes controlled by MpERF13 in oil body cell differentiation ^7^. These include Mp*MYB02,* Mp*SYP12B, MpABCG1,* and Mp*ERF13* itself, whose expression is also under the tight control of the TF it encodes ^7^. We hypothesized that if MpERF13 TF is a bona fide target of MpNPR, plants with altered levels of MpNPR, such as Mp*npr* mutants and overexpressing MpNPR lines, must show altered patterns of expression for MpERF13-dependent genes. To test this, we performed firstly quantitative RT-PCR analysis for Mp*MYB02,* Mp*SYP12B, MpABCG1,* and also Mp*ERF13*, on Mp*npr* and Mp*erf13-1^ge^* mutants grown on non-axenic conditions. Expression of all those genes was almost undetectable in the Mp*erf13-1^ge^* mutant as previously described ^7^, while both Mp*npr* alleles showed significant lower expression compared with Tak-1, indicating that MpERF13-dependent gene expression is partially compromised in plants lacking Mp*NPR* (Figure 3A). We further compared the expression of those genes in Mp*NPR^ox^* and Mp*erf13^GOF^* plants. Contrary to previous results, all those genes were highly expressed in plants overexpressing Mp*NPR* compared to Tak-1, although to a lesser extent than Mp*erf13^GOF^*, which showed remarkably high levels of expression for all genes (Figure 3B). Altogether, our results indicate that MpNPR acts as a positive regulator of MpERF13-dependent transcriptional activity, including MpERF13 itself.

**Figure 3.**
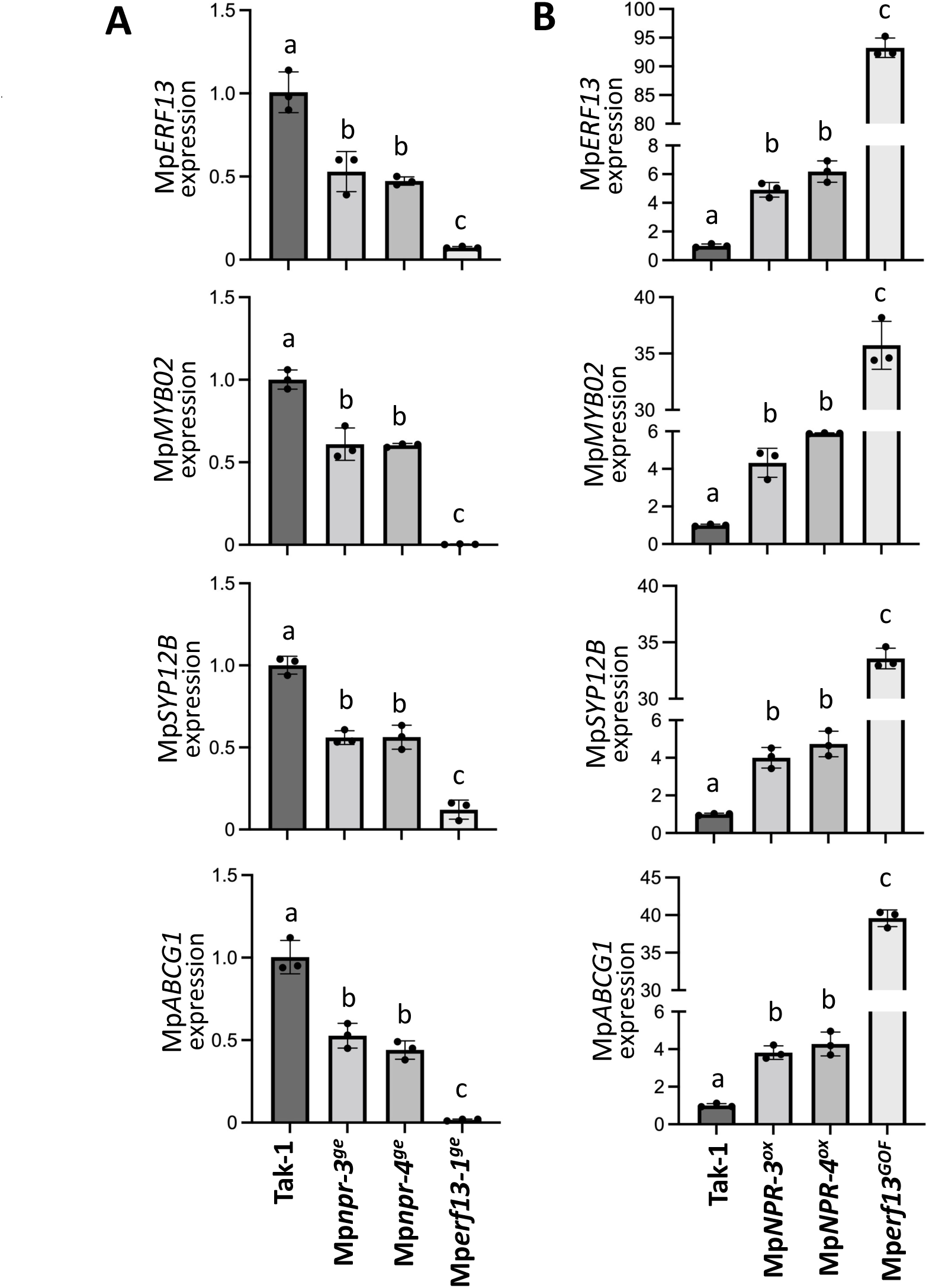
MpNPR regulates the expression of MpERF13-dependent genes. **(A)** qPCR analysis of three selected MpERF13-dependent marker genes (Mp*MYB02*, Mp*SYPB12B* and Mp*ABCG1*) in WT Tak-1, Mp*npr-3^ge^,* Mp*npr-4^ge^* and Mp*erf13-1^ge^* plants grown in vermiculite for five days (basal conditions). Gene expression was normalized against Mp*EF1⍺.* The measurements represent the relative expression compared to WT Tak-1. Error bars represent SD (n = 3; biological replicates). Letters indicate statistically significant groups (one-way ANOVA, Tukey’s HSD, *p* < 0.01). **(B)** qPCR analysis of three selected MpERF13-dependent marker genes (Mp*MYB02*, Mp*SYPB12B* and Mp*ABCG1*) in WT Tak-1, Mp*NPR-3^ox^,* Mp*NPR-4^ox^* and Mp*erf13^GOF^* plants grown in vermiculite for five days (basal conditions). Gene expression was normalized against Mp*EF1⍺.* The measurements represent the relative expression compared to WT Tak-1. Error bars represent SD (n = 3; biological replicates). Letters indicate statistically significant groups (one-way ANOVA, Tukey’s HSD, *p* < 0.05).

### MpNPR controls oil body formation through MpERF13 transcription factor

Our previous data suggest that MpNPR functions as an MpERF13 coregulator during oil body cell differentiation in *Marchantia*. Thus, we next examined whether ectopic expression of MpNPR leads to activation of Mp*ERF13,* Mp*MYB02,* Mp*SYP12B,* and *MpABCG1* genes in an MpERF13-dependent manner. To do this, we firstly generated transgenic *Marchantia* plants overexpressing Mp*NPR-citrine* under the control of the Mp*EF1⍺* promoter in the Mp*erf13-1^ge^* mutant background. Western blotting assays confirmed protein accumulation of MpNPR-citrine, and two independent lines showing similar protein accumulation to the previously generated Mp*NPR-3^ox^* and Mp*NPR-4^ox^*plants were selected (Figure S5). These lines were designated as Mp*NPR-5^ox^*Mp*erf13-1^ge^* and Mp*NPR-8^ox^* Mp*erf13-1^ge^*, respectively. Then, we performed by quantitative RT-PCR analysis for the activation of those genes in Mp*erf13-1^ge^* and Mp*NPR-4^ox^* as controls, and both Mp*NPR^ox^* Mp*erf13-1^ge^*alleles grown in non-axenic conditions. As seen before, all those genes were highly expressed in Mp*NPR-4^ox^*, but their expression was significantly lower or not detectable in the Mp*erf13-1^ge^* mutant compared with Tak-1 plants (Figure 4A). In all cases, expression levels of those genes on both Mp*NPR^ox^* Mp*erf13-1^ge^* alleles were similar to the Mp*erf13-1^ge^* mutant (Figure 4A). This indicates that MpNPR induces the activation of oil body-related genes in an MpERF13-dependent manner.

**Figure 4.**
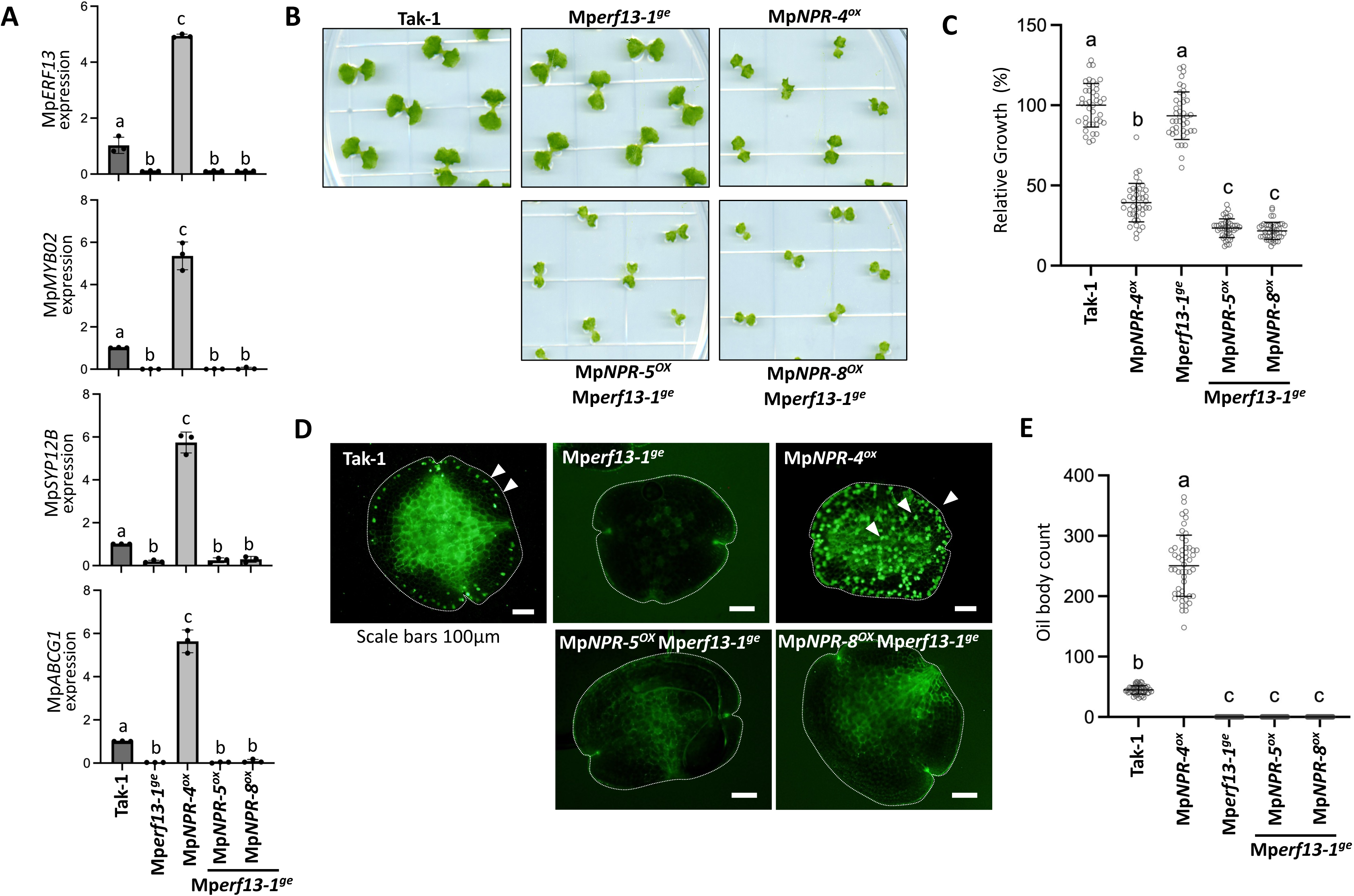
MpNPR controls oil body formation through MpERF13 transcription factor. **(A)** qPCR analysis of selected MpERF13-dependent marker genes (Mp*MYB02*, Mp*SYPB12B,* Mp*ABCG1* and Mp*ERF13*) in WT Tak-1, Mp*erf13-1^ge^*, Mp*NPR-4^ox^,* Mp*NPR-5^OX^*Mp*erf13-1^ge^* and Mp*NPR-8^OX^* Mp*erf13-1^ge^* plants grown in vermiculite for five days (basal conditions). Gene expression was normalized against Mp*EF1⍺.* The measurements represent the relative expression compared to WT Tak-1. Error bars represent SD (n = 3; biological replicates). Letters indicate statistically significant groups (one-way ANOVA, Tukey’s HSD, *p* < 0.05). **(B)** Phenotype of 12 days-old WT Tak-1, Mp*erf13-1^ge^*, Mp*NPR-4^ox^* and overexpression of Mp*NPR* in the Mp*erf13-1^ge^*mutant background (Mp*NPR-5^OX^* Mp*erf13-1^ge^* and Mp*NPR-8^OX^* Mp*erf13-1^ge^* lines). Growth inhibition induced by overexpression of Mp*NPR* in Tak-1 is not reverted in plants lacking Mp*erf13*. **(C)** Relative growth (% area) of plants from (A) compared to Tak-1. Data shown as mean ± SD (n=14). Different letters represent statistical significances (one-way ANOVA, Tukey’s HSD, *p* < 0.01). **(D)** Phenotype of BODIPY-stained 0-d-old gemmae from WT Tak-1 Mp*erf13-1^ge^*, Mp*NPR-4^ox^* and overexpression of Mp*NPR* in the Mp*erf13-1^ge^* mutant background (Mp*NPR-5^OX^* Mp*erf13-1^ge^* and Mp*NPR-8^OX^* Mp*erf13-1^ge^* lines) under fluorescence microscopy. Oil bodies are indicated by arrows. **(E)** Number of oil bodies visualized by BODIPY-staining in 0-d-old gemmae of WT Tak-1 gemmae from WT Tak-1 Mp*erf13-1^ge^*, Mp*NPR-4^ox^* and overexpression of Mp*NPR* in the Mp*erf13-1^ge^* mutant background (Mp*NPR-5^OX^* Mp*erf13-1^ge^* and Mp*NPR-8^OX^* Mp*erf13-1^ge^* lines). Bars indicate means ± SD (n = 50). Letters indicate statistically significant groups (one-way ANOVA, Tukey’s HSD, *p* < 0.01). **(E)** qPCR analysis of selected MpERF13-dependent marker genes (Mp*MYB02*, Mp*SYPB12B,* Mp*ABCG1* and Mp*ERF13*) in WT Tak-1, Mp*erf13-1^ge^*, Mp*NPR-4^ox^,* Mp*NPR-5^OX^* Mp*erf13-1^ge^*and Mp*NPR-8^OX^* Mp*erf13-1^ge^* plants grown in vermiculite for five days (basal conditions). Gene expression was normalized against Mp*EF1⍺.* The measurements represent the relative expression compared to WT Tak-1. Error bars represent SD (n = 3; biological replicates). Letters indicate statistically significant groups (one-way ANOVA, Tukey’s HSD, *p* < 0.05).

Finally, we next examined whether MpNPR-induced oil body formation also requires the MpERF13 TF. Notably, similar to the overexpression of Mp*NPR* in the Tak-1 background (Mp*NPR-4^ox^*), both Mp*NPR-5^ox^* Mp*erf13-1^ge^* and Mp*NPR-8^ox^* Mp*erf13-1^ge^* plants also showed a marked inhibition of growth compared to the Mp*erf13-1^ge^* background (Figure 4B and 4C). This indicates that ectopic expression of MpNPR in *Marchantia* leads to thallus growth inhibition in a manner that is independent of MpERF13 (Figure 4B and 4C). In contrast, gemmae from both Mp*NPR^ox^* Mp*erf13-1^ge^* alleles did not resemble plants overexpressing Mp*NPR* in the Tak-1 background, as they were completely depleted of oil bodies to a similar extent as Mp*erf13-1^ge^* gemmae (Figure 4D and 4E), supporting that this is a specific phenotype related to the function of MpERF13. These results evidence that the ability of MpNPR to induce the formation of oil body cells is fully dependent on the MpERF13 TF in *Marchantia*, while also demonstrate that NPR mediates a novel regulatory pathway in oil body formation through the MpERF13 TF in *Marchantia*.

### MpNPR and MpERF13 control defence against snail herbivory

Recent evidence has shown the protective role of oil bodies against arthropod herbivory in *Marchantia,* as mutant plants defective in oil body cell formation, such as Mp*erf13,* are more susceptible to the terrestrial isopod *Armadillidium vulgare* than WT plants ^6,7^. Noteworthy, early experiments associated the presence of oil bodies by using fresh and alcohol-washed liverworts to the feeding behaviour of snails ^18^ and recently, *Marchantia*’s defence mechanisms were shown to be active against gastropod herbivory ^38^. Thus, we decided to evaluate the protective role of oil bodies and the function of MpERF13 and MpNPR towards the attack of snails. To do this, we firstly performed concomitant two-choice experiments using Tak-1 and Mp*erf13-1^ge^* plants on one side, and Tak-1 and Mp*npr-4^ge^* mutants on the other side, challenged with juveniles of the land snail *Helix aspersa*. Then, we monitored the total thalli area consumed by the snails for 24 h. Compared to Tak-1, thalli of Mp*erf13-1^ge^* were severely consumed while Mp*npr-4^e^*mutants were significantly reduced, although to a lesser extent than Mp*erf13-1^ge^*(Figure 5A and 5B). These results correlate with the level of previously observed defects in oil body content on both Mp*erf13* and Mp*npr* mutant gemmae (Figure 2A and 2B). We next performed concomitant two-choice experiments comparing Tak-1 with Mp*erf13^GOF^*or Mp*NPR-4^ox^*, as gemmae from those plants showed in our experiments a similar enhanced pattern for oil body distribution and numbers. Thallus areas of Tak-1 plants were significantly reduced compared with Mp*erf13^GOF^*or Mp*NPR-4^ox^* thalli, which in both cases, remained almost intact, and to a similar extent after 24 h of herbivory (Figure 5C and 5D). This indicates that ectopic expression of NPR or MpERF13 in *Marchantia* leads to gastropod resistance. Finally, we evaluated whether enhanced resistance to the attack of land snails observed in Mp*NPR^ox^* plants is dependent on MpERF13. To test this, we performed four simultaneously two-choice experiments comparing Tak-1 with Mp*NPR-4^ox^*or Mp*erf13-1^ge^* as controls, or both Mp*NPR^ox^* Mp*erf13-1^ge^* generated independent alleles. As shown in Figures 5E and 5F, MpNPR-triggered enhanced resistance against snail herbivory was fully abolished when this gene was introduced into the Mp*erf13-1^ge^* mutant background (Figure 5E and 5F). Indeed, Mp*NPR^ox^* Mp*erf13-1^ge^*alleles and the Mp*erf13-1^ge^* mutant showed a similar enhanced susceptibility to the attack of snails (Figure 5E and 5F). These data demonstrate that the ability of MpNPR to induce resistance to gastropods is fully dependent on MpERF13, while also expands the protective role of oil bodies in liverworts towards the defence against the attack of gastropods. Taken together, our results indicate that MpNPR controls lineage-specific oil body formation and defence against gastropod herbivory through the MpERF13 TF in *Marchantia*.

**Figure 5.**
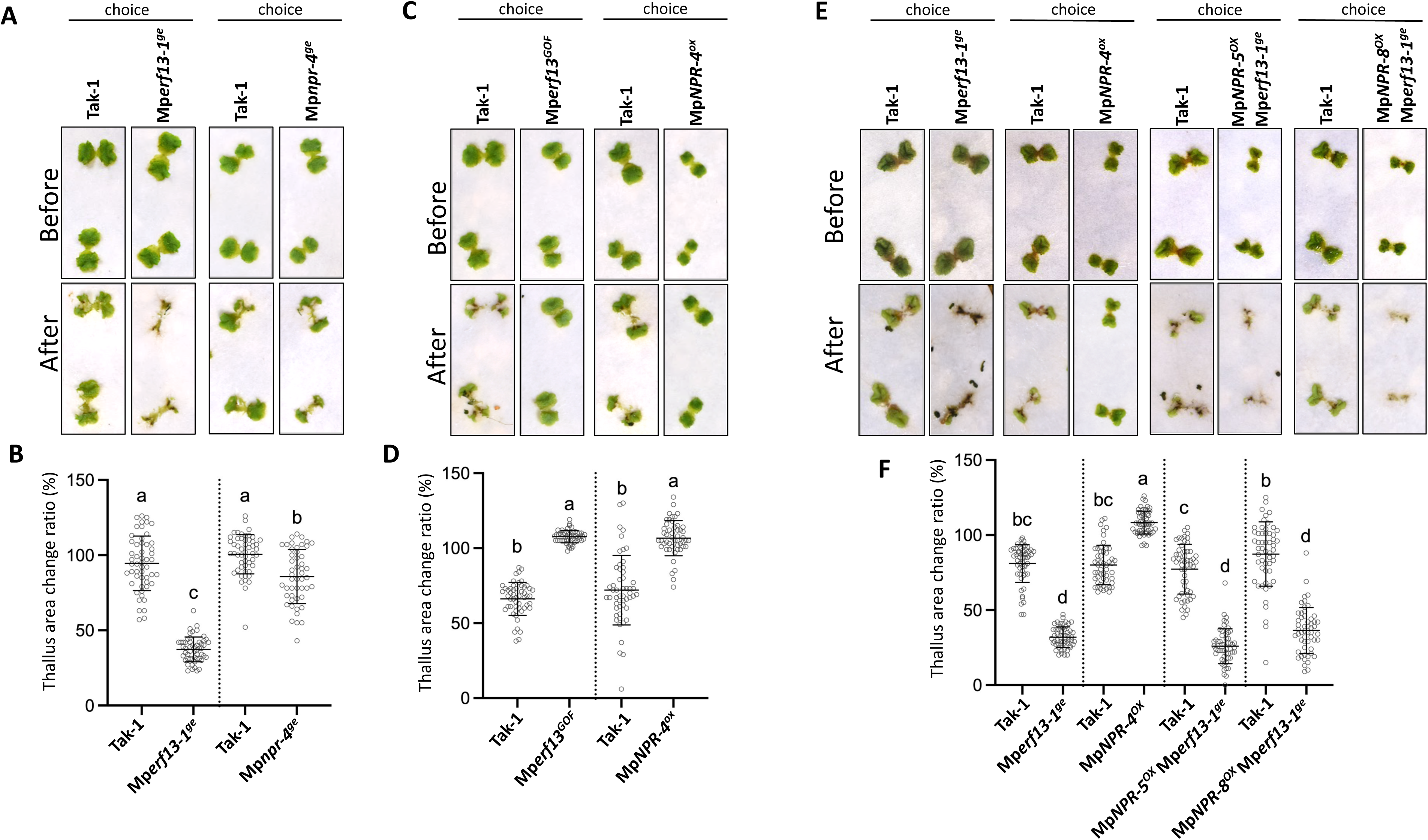
MpNPR and MpERF13 control defence against snail herbivory. **(A)** 12-day-old thalli of WT Tak-1, Mp*erf13-1^ge^* and Mp*npr-4^ge^* plants before and after co-cultivation with snails (*Helix aspersa*) for 24 hours. Herbivory experiments were designed as a two-choice experiment, comparing WT Tak-1 with Mp*erf13-1^ge^* or Mp*npr-4^ge^*. Both two-choice experiments were performed at the same time and thus, they are directly comparable. **(B)** Quantification of snail feeding assay with 12-d-old WT Tak-1, Mp*erf13-1^ge^* and Mp*npr-4^ge^* plants from (A). Plants were co-cultivated for 24 h with snails (*Helix aspersa*) and thallus area change ratio (%) was determined by area quantification after and before the feeding assay. Dot lines separate two-choice experiments performed at the same time. Bars indicate means ± SD (n = 54). Letters indicate statistically significant groups (one-way ANOVA, Tukey’s HSD, *p* < 0.01). **(C)** 12-day-old thalli of WT Tak-1, Mp*erf13^GOF^* and Mp*NPR-4^OX^* plants before and after co-cultivation with snails (*Helix aspersa*) for 24 hours. Herbivory experiments were designed as a two-choice experiment, comparing WT Tak-1 with Mp*erf13^GOF^* or Mp*NPR-4^OX^*. Both two-choice experiments were performed at the same time and thus, they are directly comparable. **(D)** Quantification of snail feeding assay with 12-d-old WT Tak-1, Mp*erf13^GOF^* and Mp*NPR-4^OX^* plants from (C). Plants were co-cultivated for 24 h with snails (*Helix aspersa*) and thallus area change ratio (%) was determined by area quantification after and before the feeding assay. Dot lines separate two-choice experiments performed at the same time. Bars indicate means ± SD (n = 54). Letters indicate statistically significant groups (one-way ANOVA, Tukey’s HSD, *p* < 0.01). **(E)** 12-day-old thalli of WT Tak-1, Mp*erf13-1^ge^,* Mp*NPR-4^OX^,* Mp*NPR-5^OX^* Mp*erf13-1^ge^* and Mp*NPR-8^OX^*Mp*erf13-1^ge^* plants before and after co-cultivation with snails (*Helix aspersa*) for 24 hours. Herbivory experiments were performed comparing WT Tak-1 with Mp*erf13-1^ge^,* Mp*NPR-4^OX^,* Mp*NPR-5^OX^* Mp*erf13-1^ge^* or Mp*NPR-8^OX^* Mp*erf13-1^ge^* plants, as four two-choice experiments performed at the same time and thus, they are comparable. **(F)** Quantification of snail feeding assay with 12-d-old WT Tak-1, Mp*erf13-1^ge^,* Mp*NPR-4^OX^,* Mp*NPR-5^OX^* Mp*erf13-1^ge^* and Mp*NPR-8^OX^* Mp*erf13-1^ge^* plants from (E). Plants were co-cultivated for 24 h with snails (*Helix aspersa*) and thallus area change ratio (%) was determined by area quantification after and before the feeding assay. Dot lines separate two-choice experiments performed at the same time. Bars indicate means ± SD (n = 54). Letters indicate statistically significant groups (one-way ANOVA, Tukey’s HSD, *p* < 0.01).

## DISCUSSION

Since its discovery more than 20 years ago, NPR proteins have been the focus of countless studies in a multitude of vascular plants and agricultural crops ^24^. In *Arabidopsis*, NPR genes play essential roles as hormonal SA sensors and executors of SA-dependent plant defences against pathogens with prominent biotrophic or hemibiotrophic lifestyles such as *P. syringae*. But surprisingly, the recent characterization of the unique MpNPR has not supported a similar function in *Marchantia* as in angiosperms ^25^. As a paradox, while Mp*NPR* can complement the At*npr1* mutant for all tested responses so far when ectopically expressed in *Arabidopsis,* mutations in Mp*NPR* do not result in At*npr1*-like phenotypes in *Marchantia,* indicating that Mp*NPR* may not be the master regulator of SA-induced responses in this liverwort ^25,26^. Indeed, hornworts accumulate SA but lack NPR genes ^28–30^, which opens the intriguing possibility that additional SA sensors might exist in bryophytes, as it also happens in angiosperms ^39^. Nevertheless, all these compelling data suggest that NPR-associated pathways might have evolved distinctly in divergent land plant lineages ^2,25^. Here, we discover that MpNPR controls the formation of oil bodies, a synapomorphy of liverworts that function as active sites for the biosynthesis and storage of diverse bioactive specialized metabolites with cytotoxic activity ^8^. We found that genome-edited mutants of Mp*NPR* show a reduced number of oil bodies, while its overexpression in *Marchantia* induces a remarkable overaccumulation of these organelles in gemmae. And several lines of evidence support that MpNPR acts through the master MpERF13 TF for controlling the signalling pathway leading to oil body cell differentiation in this liverwort. Firstly, MpNPR interacts with MpERF13. Secondly, Mp*npr* mutants and transgenic lines overexpressing MpNPR display alterations in the expression pattern of genes that are under the control of the MpERF13 TF. These include Mp*SYBP12B*, Mp*MYB02,* and Mp*ABCG1*, all of which are responsible for critical and different steps in the formation and functionality of this organelle ^7,12,15,17,40^. Notably, Mp*ERF13* expression is also affected in plants with altered levels of Mp*NPR*, which goes in line with previous observations indicating that MpERF13 is also under its own transcriptional control, which further supports the close regulation of MpERF13 by MpNPR ^7^. And finally, the effect of MpNPR in regulating oil body formation, is fully dependent on MpERF13. Globally, our results demonstrate that NPR mediates a novel regulatory pathway in lineage-specific oil body formation through the MpERF13 TF in *Marchantia*.

Notably, despite the overexpression of Mp*NPR* in *Marchantia* leads to the overaccumulation of oil bodies in gemmae similar to that observed for the gain-of-function Mp*erf13^GOF^*, Mp*npr* mutants do not resemble to Mp*erf13* plants, which are completely depleted of oil bodies. Indeed, Mp*npr* plants only display subtle defects in oil body cell counts, that are evident when plants are grown under non-axenic conditions, but not on axenic conditions. While this likely explains why a previous work failed to detect defects in oil body numbers on their axenically grown Mp*npr* plants compared to wild-type *Marchantia* ^16^, it also indicates that MpNPR is mostly relevant during stress-induced modulation of oil body development, likely triggered by non-axenic growth. Nonetheless, our results also indicate that MpNPR is not the unique protein modulating oil body formation in *Marchantia.* Indeed, our results suggest that other pathway/s might contribute redundantly with MpNPR towards oil body differentiation. In this sense, the OPDA signalling pathway in *Marchantia*, which is analogous to Jasmonic Acid (JA) in tracheophytes, emerges as a plausible candidate. Indeed, this signalling pathway in *Marchantia* has been widely related to the resistance against herbivory in this liverwort ^38,41,42^. Moreover, mutation in the unique Mp*MYCX/Y* TF responsible for all OPDA responses is defective in the accumulation of sesquiterpenes such as thujopsene, β-chamigrene, and cuparene, which are present in oil bodies, in response *Spodoptera littoralis* feeding ^41^. Notably, it was recently reported that the *Marchantia* OPDA receptor MpCOI1 is required for resistance against snail herbivory ^38^. All these data open the possibility that OPDA though its receptor MpCOI1 and its co-receptor MpJAZ ^42,43^, might hint additional TF targets such as MpERF13, MpC1HDZ or MpMYB02 apart from its canonical Mp*MYCX/Y* TF to modulate other responses such as oil body cell differentiation in *Marchantia.* However, as mutants in Mp*MYCX/Y* TF already contain defects on the levels of secondary metabolites accumulating in oil bodies, the most likely possibility is that OPDA might regulate any of the previously described oil-bodies related TFs at the transcriptional level through Mp*MYCX/Y*, although this remains to be demonstrated. Alternatively, abscisic acid (ABA) signalling has been shown to induce the biosynthesis of bisbibenzyls, also characteristic metabolites found in oil bodies, upon UV-C light stress ^44^. Whether any of these pathways are involved in oil body development as a mechanism to provide inbuilt redundancy with MpNPR is still unknown, and will deserve further investigations.

Finally, emerging data indicate that the oil body likely represents a predominant liverwort-specific immune innovation, as plants with reduced number of these organelles, such as mutants for the Mp*erf13* and Mp*c1hdz* TFs, has shown to exhibit increased isopod herbivory to the common pill-bug *Armadillidium vulgare*, but not defects associated to multiple abiotic stress responses in *Marchantia* ^6,7^. Moreover, the metabolic content of oil bodies itself, full of diverse biologically active cytotoxic compounds, also supports a major role of this organelle as a defensive organ against pathogen attack ^6,8,10–13^. Interestingly, extracts from plants with reduced contents of specific compounds accumulating in oil bodies have reduced antimicrobial activity against bacteria and fungi *in vitro* ^6,45^. However, whether the oil body itself is important for defence against these types of microbes still remains to be demonstrated, as it is unclear how metabolic compounds contained in these intracellular organelles can interact with these microorganisms during plant-microbe interactions in nature. Nonetheless, currently there is convincing evidence for the immune role of oil bodies against arthropod herbivory in *Marchantia* ^6,7^. Our work now expands the protective role of oil body organelles towards gastropod herbivory, as loss-of-function Mp*npr* and Mp*erf13* mutants with reduced content of oil bodies are more susceptible to land snails, while *Marchantia* plants over accumulating these organelles through the ectopic expression of MpNPR or a gain-of-function allele of Mp*ERF13* are in both cases highly resistant to the attack of this gastropod. Moreover, we found that the ability of MpNPR to protect *Marchantia* against snail feeding is directly related to the MpERF13 function, as plants overexpressing Mp*NPR* in the Mp*erf13* mutant background are highly susceptible to snail feeding to a similar extent as Mp*erf13^ge^*mutants. On the one side, our results provide now convincing evidence supporting the hypothesis formulated more than a century ago, that oil bodies from liverworts contain deterrent compounds that are effective against land snails ^18^. And on the other side, we uncover the existence of an Mp*NPR-*Mp*ERF13* module controlling oil body differentiation and defences against snail herbivory. Our results are appealing as in angiosperms NPRs have profoundly associated to the resistance against biotrophic or hemibiotrophic pathogens such as *P. syringae*. However, a similar scenario is currently not supported in *Marchantia*. Indeed, Mp*npr* mutants were recently found to be more resistant to hemibiotrophic bacteria and hypersensitive to exogenous SA treatments, suggesting that indeed, MpNPR might function as a rather negative regulator for these responses ^25^. While the molecular mechanisms for those outcomes still remain to be uncovered, it is possible that they might reflect additional specialised trajectories for the unique MpNPR in *Marchantia* to defend against pathogens, as it has been previously proposed ^2,25^. NPR proteins originated in the most recent common ancestor of land plants, during the early stages of terrestrialization ^26^. Taking into account that the estimated origin of gastropods in the Cambrian–Ordovician era coincides with that of the first land plants ^38^, and that several fossil evidences suggest early herbivory on ancestral liverworts ^46,47^, it is tempting to speculate that NPRs from liverworts were co-opted towards the control of oil body, as a unique lineage-specific immune strategy, to fend off against the specific selective pressures that early herbivores impose on ancestral liverworts. Moreover, as oil bodies are a synapomorphy of liverworts, our results pinpoint an unique specialised evolutionary trajectory of NPR in liverworts.

## RESOURCE AVAILABILITY

### Lead Contact

Requests for new resources generated in this study and further information should be directed toward Selena Gimenez-Ibanez.

### Materials Availability

New plasmids and plant materials generated in this study are available upon request. Please note that the distribution of generated transgenic plants will be governed by material transfer agreements (MTAs) and will be dependent on appropriate import permits acquired by the receiver.

### Data and Code Availability

No datasets has been generated during this study.

## ACKNOWLEDGEMENTS

The authors are grateful to the Department of “Advanced Optical Microscopy” at CNB-CSIC for technical assistance in the confocal analyses of MpNPR-citrine. We thank Prof. T. Ueda for the Mp*erf13-1^ge^* and Mp*erf13^GOF^* mutant plants. We thank Jose Antonio Marcelo from Cargols de Cal Jep for providing juvenile common land snails (*Helix aspersa*). S.M. was funded by a PRE2020-093780 grant by MCIN/AEI/10.13039/501100011033. R.S. was funded by a PID2022-140766OB-I00 grant by the MCIN/AEI/10.13039/501100011033 and ‘‘ERDF/EU’’. This work was funded by the PID2022-136746OB-I00 Grant by the MCIN/AEI/10.13039/501100011033 and ‘‘ERDF/EU’’ to S.G.-I.

## AUTHOR CONTRIBUTIONS

S.G-I. designed the research; S.G-I. conceived the project; L.E-C. and S.M. generated materials; L.E-C., S.M. and M.G-Z. performed experiments; L.E-C. and S.G-I analysed the data; R.S. and S.G-I. discussed the data; L.E-C., R.S. and S.G-I. wrote the manuscript; All authors corrected the manuscript.

## DECLARATION OF INTERESTS

The authors declare no competing interests.

## MATERIAL AND METHODS

### Plant material

*Marchantia polymorpha* accession Takaragaike-1 (Tak-1; male) was used as wild-type (WT). To generate Mp*npr* mutants in the Tak-1 background, we used CRISPR-Cas9 nickase-mediated mutagenesis targeting Mp*NPR* (Mp1g02380). Four different gRNAs were designed to target the gene (listed below). These gRNAs were first cloned into the vectors pMPGE_EN04, pBC-GE12, pBC-GE23, and pBC-GE34. Subsequently, they were transferred into the pMpGE018 binary vector, which carries the CRISPRCas9 nickase, using Gateway LR Clonase (Invitrogen). Tak-1 plants were transformed, and thalli were selected by chlorosulfuron resistance employing the regenerating thalli transformation method ^48^. To identify mutant plants, genomic DNA of chlorosulfuron-resistant explants was extracted and sequenced using primers flanking the gRNAs target sites. For the generation of Mp*NPR-citrine* overexpression in Tak-1 or Mp*erf13-1^ge^* backgrounds, the full-length CDS of Mp*NPR* (Mp1g02380), designed with Gateway attB sites and lacking the stop codon, was synthesized by Integrated DNA Technolgies (IDT, Coralville, Iowa, USA), cloned into the pDONR207 vector (Invitrogen) and then, transferred into the binary vector pMpGWB108 ^49^, which express Mp*NPR* fused to *citrine* under the control of the Mp*EF1α* promoter (*pro*Mp*EF1α:*Mp*NPR-citrine*). Tak-1 or Mp*erf13-1^ge^* plants were transformed and selected by hygromycin resistance as previously described. Mp*erf13-1^ge^* and Mp*erf13^GOF^* were previously described, and kindly obtained by Prof. T. Ueda (NIBB) (Kanazawa et al., 2020).

### Gene identification

Sequences were obtained from the Plant Genomics Resource Phytozome (https://phytozome-next.jgi.doe.gov) and Genome Database for Marchantia (https://marchantia.info).

### Growth conditions

*Marchantia* gemmae were grown on half Gamborg’s B5 medium containing 1% agar, under long day conditions (16 h light; 50–60 µmol m−2 s−1) *in vitro* at 22 °C, for 5 to 10 days for all experiments to synchronize an homogenous initial growth of all gemmae, commonly on whatman filter papers. Then, plants were transferred to non-axenic conditions, on pots or plates containing vermiculite, and grown in controlled environment chambers at an average temperature of 22°C (range 19°C–23°C) with 45%–65% relative humidity under long-day conditions (16 h light), for the specified days described for each experiment.

### Confocal Microscopy

For subcellular localization of MpNPR, gemmalings of Mp*NPR^ox^* (*pro*Mp*EF1α:*Mp*TGA-citrine/*Tak-1) lines were imaged using the confocal laser scanning microscopes STELLARIS 5 from Leica with a water-immersion objective (HC PL APO CS2 63x/1.29). Excitation and emission were set at 488 and 520 nm, respectively, for Citrine. Chloroplasts fluorescence was detected between 659 and 735 nm. Confocal images were processed using the ImageJ software ^50^.

### Fluorescence microscopy and oil body quantification

For oil body quantification, plants were grown on pots with vermiculite under non-axenic conditions, until plants generated gemma cups. Similar gemma cups were selected in all plants, and gemmae was obtained from a pool of about 5-10 gemma cups. For oil body quantification, 0-day-old gemmae were stained by 4,4-difluoro-1,3,5,7,8-pentamethyl-4-bora-3a,4a-diaza-s-indacene (BODIPY 493/503, Thermo Fisher) as previously described ^7^. Gemmae were imaged using ×5 (numerical aperture 0.12) and ×10 (numerical aperture = 0.25) objectives on a Leica DM R fluorescence microscope equipped with an MPS60 camera (Leica). Images were acquired with a GFP3 excitation filter and processed using the ImageJ software ^50^.

### Snail feeding assays

For snail feeding assays, two-choice experiments were designed by growing gemmae from *Marchantia* WT Tak-1 and the selected line to be analysed on the same whatman filter paper on previously described *in vitro* conditions for 6 to 8 days. Then, filter papers containing plants were transferred to plates with vermiculite under non-axenic conditions for additional 5 to 7 days. Juvenile common land snails (*Helix aspersa*) were kindly obtained from “Cargols de Cal Jep” (https://www.caljep.com). Snails were maintained on plastic garden boxes containing vermiculite moistened with water in controlled environment chambers under long-day conditions (16 h light and 22 °C). Snails were fed with sweet potato and kept in a moist environment by water spraying once a week until use, as previously described ^38^. Two-choice experiments were conducted by placing 10 juvenile snails (*Helix aspersa*) into each plate. To quantify the area of plant tissue consumed, photos were taken before and 24 h after feeding. Thallus areas were measured using the ImageJ software ^50^, and the thallus area change ratio (%) was determined by area quantification after and before the feeding assay.

### Quantitative RT-PCR

For oil body quantification, plants were grown on pots with vermiculite under non-axenic conditions for 5-7 days. RNA was extracted and quantitative RT-PCR was performed as previously described ^51^. Data analysis shown was done using three biological replicates. Error bars represent standard deviation (SD). All samples were normalized against the housekeeping gene Mp*EF1α* (Mp3g23400).

### Yeast-two hybrid

For yeast two-hybrid assays, the full-length CDS of Mp*ERF13* (Mp6g08690), designed with Gateway attB sites and lacking the stop codon, was synthesized by Integrated DNA Technolgies (IDT, Coralville, Iowa, USA). The gene was cloned into pDONR207 vector with a Gateway BP Clonase II kit (Invitrogen) and verified by sequencing. Both Mp*ERF13* and Mp*NPR* were then transferred into the pGADT7 (AD-Gal4) and Gal4-based pGBKT7 (BD-Gal4) vectors (Clontech Laboratories Inc.), respectively, using the Gateway system (Invitrogen). The bait and prey plasmids were transformed into *S. cerevisiae* strain AH109, and transformants were grown on plasmid-selective media (SD/-Trp-Leu). Plates were incubated at 28°C for 4 days and independent colonies for each bait-prey combination were resuspended in sterile water. Five microliters of each suspension were spotted onto two alternative interaction-selective media (SD/-Trp-Leu-Ade-His, and SD/-Trp-Leu-Ade-His + 10mM 3-amino-1,2,4-triazole). Plates were incubated at 28°C and photographed after 4 or 7 days of incubation.

### Protein extract and western blotting

For detection of the MpNPR-citrine, total proteins from Marchantia Tak-1 and Mp*NPR^OX^* (*pro*Mp*EF1a:*Mp*TGA-citrine/*Tak-1) plants were extracted in extraction buffer (50 mM Tris-HCl pH 7.5, 150 mM NaCl, 1 mM DTT, 1 mM PMSF, 10% Glycerol, protease inhibitors cocktail EDTA-free (Roche), 2mM EDTA, and 1 % Triton X-100). Protein extracts were sonicated (5 cycles 15 sec on / 15 sec off at 4°C) and then centrifuged twice at 20,000 g for 10 min at 4°C. Western blots were performed with anti-GFP-horseradish peroxidase-conjugated antibody (Miltenyi Biotec).

### Pull-down assays

For MBP-MpERF13 fusion protein, MpERF13 was transferred from pDONR207 into pKM596 by recombination (Gateway, Invitrogen) ^52^. Recombinant protein purification from *E. coli* was performed as described ^53^. Total proteins from 12-days-old *Marchantia* Tak-1 and Mp*NPR^OX^* (*pro*Mp*EF1a:*Mp*TGA-citrine/*Tak-1) plants were extracted as described here before, and used for pull-down assays as described ^53^.

### Statistical methods

The data generated were statistically analysed using GraphPad Prism software. Two-tailed Student’s t-test was used to identify statistical differences between two means. For comparisons of more than two groups, statistical confidence was established using One-way ANOVA followed by Tukey’s HSD correction for multiple comparisons.

### List of primers used in this work

CRISPR-Cas9:

NPR-gRNA1A: CTCGCAAGATCTACTCCTAATCTC

NPR-gRNA1B: AAACGAGATTAGGAGTAGATCTTG

NPR-gRNA2A: CTCGGATTTCACAATCACAGTGCA

NPR-gRNA2B: AAACTGCACTGTGATTGTGAAATC

NPR-gRNA3A: CTCGATACTCGTCAGATATTCTCC

NPR-gRNA3B: AAACGGAGAATATCTGACGAGTAT

NPR-gRNA4A: CTCGATGAAAGTCGCGAAGACTCA

NPR-gRNA4B: AAACTGAGTCTTCGCGACTTTCAT

Genotyping:

MpNPR1_3’intron1_R: CATAGCACTGGGGAAAGGAA

MpNPR1_5’intron1_F: TGACATGAGCTGGTTTGAGC

Quantitative RT-PCR:

Mp1g02380 MpNPR_F: CCTGGAGAATATCTGACGAGTA

Mp1g02380 MpNPR_R: GTCTTCCTAAGCTCCTTCACTT

Mp6g08690 MpERF13_ F: TCGGAATCTGATCAGGCCTC

Mp6g08690 MpERF13_ R: CTGAGGTTTTCTACGAGCGC

Mp4g20670 MpSYP12B_F: CTTCATCCAGAGAGCCATAC

Mp4g20670 MpSYP12B_R: TTTGGTGTAGGAATCTGCTC

Mp3g07510 MpMYB02_F: CCAGAAGATGTTGTTCAACG

Mp3g07510 MpMYB02_R: CAGAGAGTTGCTGGGTTATT

Mp8g13070 MpABCG1_F: TGAGATTCTTTCCCGCATGG

Mp8g13070 MpABCG1_R: GTCAGCGATCCCAATGAGTT

Mp3g23400 Mp*EF1⍺_*F: CCGAGATCCTGACCAAGG

Mp3g23400 Mp*EF1⍺_*R: GAGGTGGGTACTCAGCGAAG

## SUPPLEMENTAL FIGURE LEGENDS

**Supplemental Figure 1. Development of mutants for Mp*NPR* in *Marchantia* by CRISPR/Cas9 technology.**

**(A)** Mutants generated by CRISPR-Cas9^D10A^ nickase-mediated mutagenesis. Schematic diagram of the Mp1g02380 WT Tak-1 locus and mutant alleles. Light grey boxes represent exons. Conserved domain of NPR proteins is shown as a green box (BTB/POZ domain) and a pink box (the ankyrin repeat-containing region). Mp*npr*-*3^ge^* contains a deletion of 22 bp while Mp*npr*-*4^ge^* contains an insertion of 37 bp in the first exon of Mp*NPR,* that in both cases generate premature stop codons. Deletions are indicated by red dashes and insertions by blue lowercase letters.

**(B)** 12-days-old Tak-1 and Mp*npr* mutants (Mp*npr-3^ge^*and Mp*npr-4^ge^*), showing a normal developmental phenotype.

**Supplemental Figure 2. Oil body formation WT Tak-1 grown under different conditions.**

Number of oil bodies in 0-d-old gemmae of WT Tak-1 plants grown under *in vitro* conditions (1/5 GB5 medium) or vermiculite. Bars indicate means ± SD (n = 60). Asterisk indicate statistically significant differences based on a two-tailed Student’s *t*-test. \**, p* < 0.001.

**Supplemental Figure 3. Oil body formation in Mp*npr* mutants grown on agar plates (1/2 GB5 medium) under *in vitro* conditions.**

**(A)** Phenotype of BODIPY-stained 0-d-old gemmae from WT Tak-1 and Mp*npr* mutant plants under fluorescence microscopy grown on agar plates (1/2 GB5 medium) under *in vitro* conditions. Oil bodies are indicated by arrows.

**(B)** Number of oil bodies visualized by BODIPY-staining in 0-d-old gemmae of WT Tak-1 and Mp*npr* mutant plants grown on agar plates (1/2 GB5 medium) under *in vitro* conditions. Bars indicate means ± SD (n = 50). Letters indicate statistically significant groups (one-way ANOVA, Tukey’s HSD, *p* < 0.01).

**Supplemental Figure 4. Development of Marchantia plants overexpressing Mp*NPR*-*citrine*.**

**(A)** Immunoblots showing MpNPR accumulation in developed independent lines of *pro*Mp*EF1α:*Mp*NPR-citrine* in Tak-1 or WT Tak-1 control plants.

**(B)** Phenotype of 12- and 25-days-old WT Tak-1 and Mp*NPR* overexpression plants, which show a delayed growth rate.

**(C)** Subcellular localization of MpNPR in Mp*NPR-4^ox^* (*pro*Mp*EF1α:*Mp*NPR-citrine/*Tak-1) plants by confocal microscopy. Scale bar 5 μm.

**Supplemental Figure 5. MpNPR accumulation in developed independent lines of *pro*Mp*EF1α:*Mp*NPR-citrine* in the Tak-1 and Mp*erf13-1^ge^* backgrounds.**

Immunoblots showing MpNPR accumulation in developed independent lines of *pro*Mp*EF1α:*Mp*NPR-citrine* in the Tak-1 and Mp*erf13-1^ge^* backgrounds. Independent lines with similar levels of MpNPR accumulation in both Tak-1 and Mp*erf13-1^ge^* backgrounds were selected for further studies.

## Notes

### Competing Interest Statement

The authors have declared no competing interest.

