## Supplementary figures and images for "MpNPR modulates lineage-specific oil body development and defence against gastropod herbivory in *Marchantia polymorpha*"

### Supplemental Figures

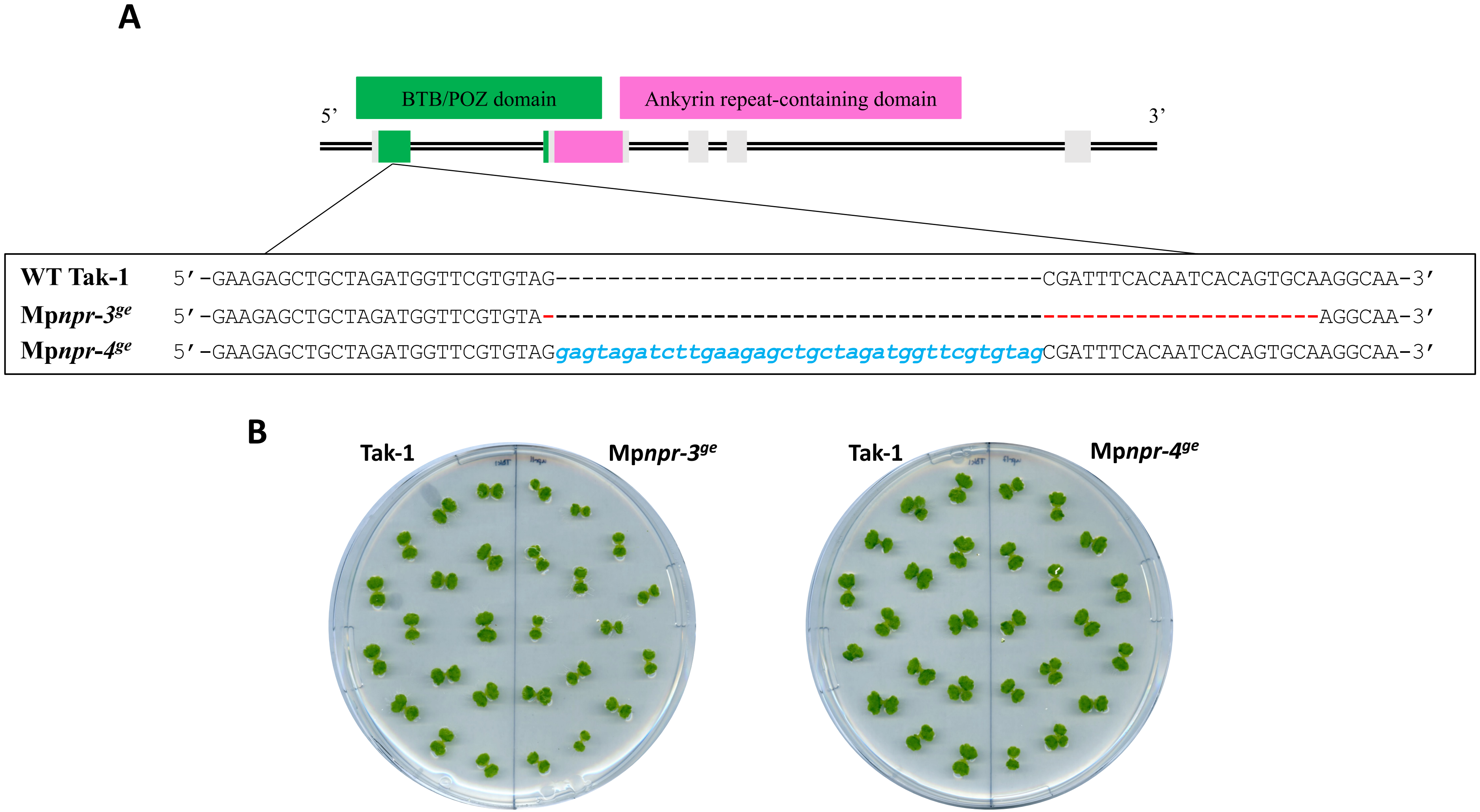

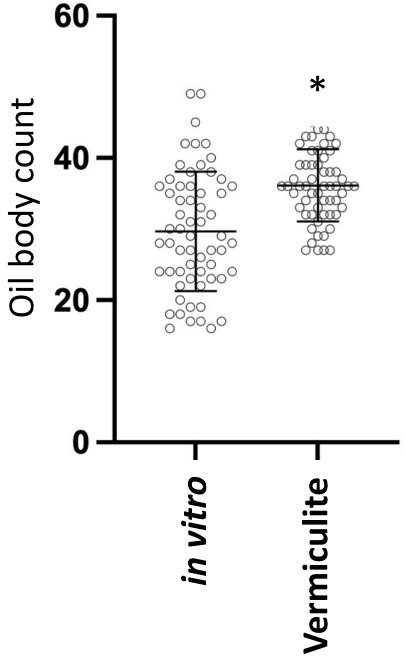


**Figure S2**

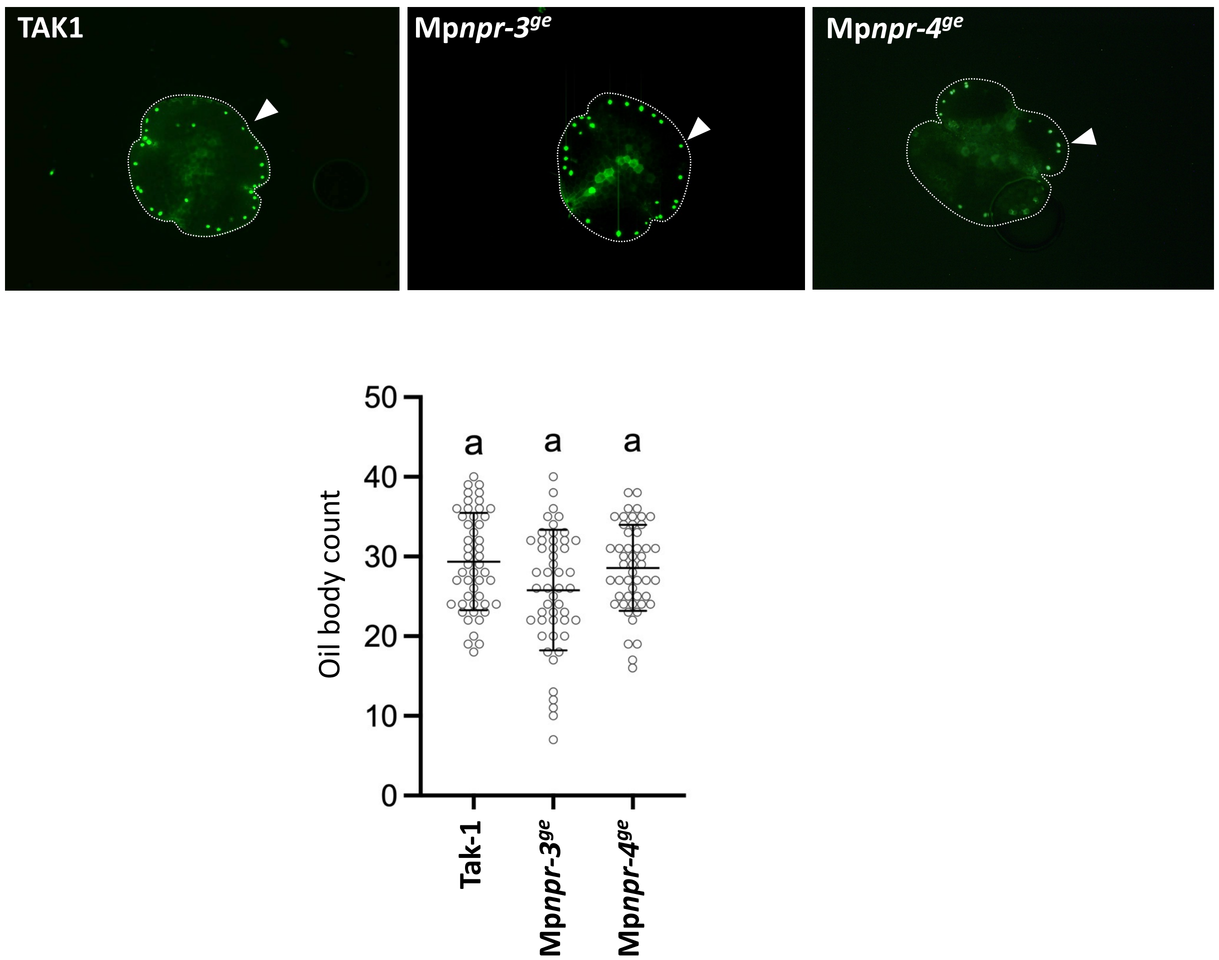

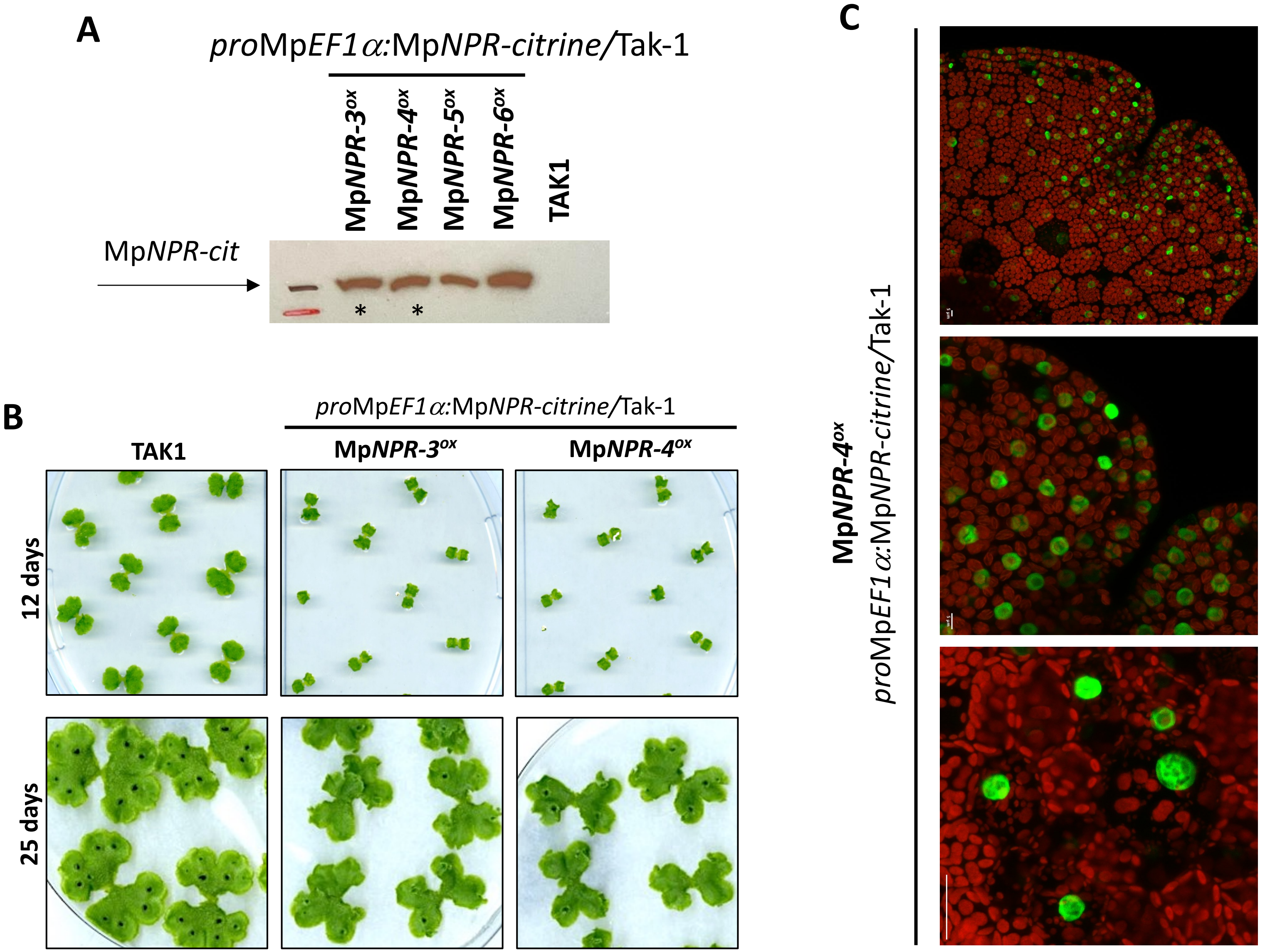

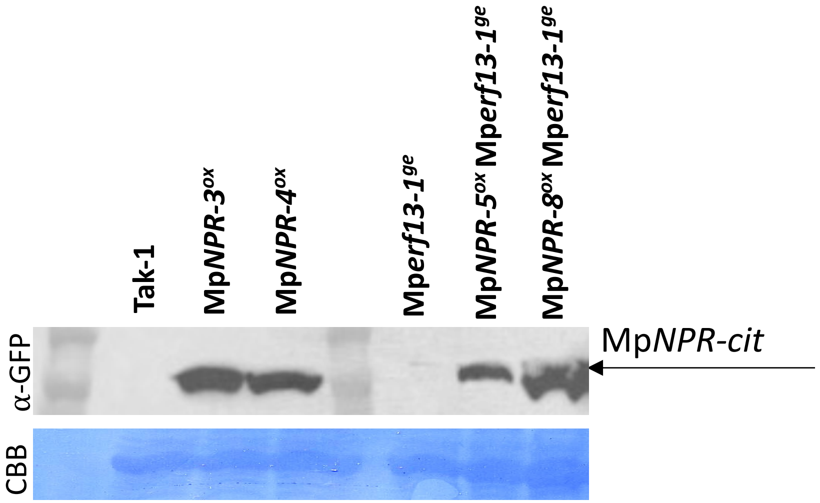
